# Autocrine inhibition of cell motility can drive epithelial branching morphogenesis in absence of growth

**DOI:** 10.1101/2020.05.15.088377

**Authors:** Elisabeth G. Rens, Mathé T. Zeegers, Iraes Rabbers, Andras Szabó, Roeland M.H. Merks

**Affiliations:** Centrum Wiskunde & Informatica, Amsterdam, The Netherlands; Leiden University, Mathematical Institute, Leiden, The Netherlands; Leiden University, Institute of Biology, Leiden, The Netherlands; Mathematics Department, University of British Columbia, Vancouver, Canada; Laboratory of Biochemistry, Wageningen University & Research, Wageningen, The Netherlands

## Abstract

Epithelial branching morphogenesis drives the development of organs such as the lung, salivary gland, kidney and the mammary gland. It involves cell proliferation, cell differentiation and cell migration. An elaborate network of chemical and mechanical signals between the epithelium and the surrounding mesenchymal tissues regulates the formation and growth of branching organs. Surprisingly, when cultured in isolation from mesenchymal tissues, many epithelial tissues retain the ability to exhibit branching morphogenesis even in absence of proliferation. In this work, we propose a simple, experimentally-plausible mechanism that can drive branching morphogenesis in absence of proliferation and cross-talk with the surrounding mesenchymal tissue. The assumptions of our mathematical model derive from *in vitro* observations of the behavior of mammary epithelial cells. These data show that autocrine secretion of the growth factor TGF*β*1 inhibits the formation of cell protrusions, leading to curvature dependent inhibition of sprouting. Our hybrid cellular Potts and partial-differential equation model correctly reproduces the experimentally observed tissue-geometry dependent determination of the sites of branching, and it suffices for the formation of self-avoiding branching structures in absence and also in presence of cell proliferation.

## 1 Introduction

During the development of organs such as lungs, kidneys and the mammary gland, epithelial tissues undergo shape changes during embryonic development resulting in a tree-like structure of branches [7, 26]. The function of branched organs is to optimize the exchange of chemicals with the surrounding tissue by maximizing the interfacial area. The dynamics of branching from an initially tube shaped epithelial tissue, called the duct, into the surrounding mesenchymal tissue involves cellular mechanisms such as directed cell migration, oriented cell division, cell shape changes, cell differentiation and cell competition (see reviews [47, 61, 65]). The specific process of branching morphogenesis varies per organ, but the key mechanisms are believed to be conserved [8, 26]. Although the dynamics of branching in various organs have been characterized well (see for instance, lung [40], kidney [66], mammary gland [15], pancreas [62]), it is still poorly understood what mechanisms drive branching morphogenesis, and which of these mechanisms are fundamental and which ones act on top of the fundamental mechanisms for ‘fine tuning’.

Mathematical modeling is a helpful tool to analyze branching morphogenesis. A first class of models asks how biological rules operating on single branches and branch tips can lead to an observed branching pattern. For example, to explain the anatomy of the urinary collecting duct tree, Davies *et al.* [9] have proposed that the ureteric tubules secrete a hypothetical repulsive factor. The tips of tubules grow towards towards lower, local concentrations of horrid at a speed inversely proportional to the concentration of the repulsive factor. Tubule tips bifurcate once the concentration of the repulsive factor drops below a threshold. The model was used to help explain observed repulsive branch interactions in explants of the collecting urinary duct trees of the mouse kidney, and to plan follow-up experiments. Scheele et al. [56] analysed the morphogenesis of the murine mammary gland using a statistical branching model. They constructed trees of which the branches bifurcate or terminate with a near equal probability. This growth process accurately reproduced the distribution of the number of branching levels in murine mammary glands and the kidney, in support of the potential homology of epithelial branching processes [56]. In a spatially extended variant of this model, growing branches were assumed to terminate as soon as they approached an existing duct, possibly due to TGF-*β* signaling [21]. This model was able to reproduce observed tissue architecture, such as local densities of branches and directional biases of branch growth. An additional rule stating that approaching branches were repelled by adjacent branches produced better fits with the observed branch density in the kidney.

A second class of mathematical models, which includes the one proposed in this paper, focuses on the cellular and molecular mechanisms responsible for branch tip initiation, branch progression, tip splitting and tip termination. For a long time, it was thought that localized, differential cell proliferation is the main driving factor of branching, but this may not always be true [61]. In the chicken and mouse lung, the buds that initiate new branches form prior to the first appearance of cell proliferation [46, 26]. Signaling factors from the mesenchyme have been proposed to drive branching [2]. However, the mesenchyme is not required either, as epithelial tissues can branch in the absence of a surrounding mesenchyme *in vitro* [52, 11, 45, 19]. In conclusion, it is still poorly understood how epithelial tissues branch autonomously in the absence of cell proliferation and the mesenchyme. Here we propose a cellular mechanism for such autonomous branching of epithelial tissues.

Epithelial branching has been proposed to be analogous to Laplacian growth, a process that underlies branching in many non-biological systems, including crystal growth [3] and viscous fingering [33]. In a mathematical modeling study of lung morphogenesis it was proposed that the epithelium branches into the surrounding mesenchyme if the mesenchyme is less viscous than the luminal fluid in the epithelium [32]. In such Laplacian growth processes, the interface of a domain advances with a velocity proportional to the gradient of a field that obeys the Laplacian equation (∇^2^*u* = 0), *i.e.* a field dominated by the diffusion equation aka heat equation [12] with *u* = 0 at the interface. Thus points of the morphology located at an interface of positive curvature, which may arise from random deviations from a initially homogeneous boundary, experience a higher gradient of the Laplacian field and will advance faster than the points at flat or concave locations of the interface. This effect is known as the Mullins-Sekerka instability. Instead of pressure and viscosity fields, Laplacian growth dynamics of tissues could also be governed by molecular concentration fields. For example, in the context of tumor branching it was proposed that cell proliferation depends on the availability of oxygen [14]. Analogously, in a Laplacian growth model of epithelial branching [22] it was proposed that growth is proportional to the local flux of fibroblastic growth factor (FGF)

Other mathematical models have studied in detail how the regulatory interactions between the epithelium and mesenchyme can drive branching. Such epithelial-mesenchymal cross-talk may regulate the highly stereotypic branching patterns of the lung [40] and the kidney [57]. Hirashima and Iwasa [24] studied epithelial-mesenchymal cross-talk in a mathematical model based on the cellular Potts model. They assumed that a deformable epithelial layer is chemoattracted to localized sources of growth factor such as FGF10 or GDNF. This chemotactic mechanism together with cell proliferation produced branches through a buckling mechanism, where the number of branches depended on the ratio between the proliferation rate and the chemotaxis speed [24]. In a further paper, Hirashima *et al.* [25] showed how secretion of SHH by progressing buds can regulate the required localized expression of FGF10 in the mesenchyme. Inhibition of FGF10 expression at high concentration of SHH, combined with activation of SHH expression at lower concentrations of SHH produce peaks of FGF10 at small distance for the bud tip. As the tip approached the lung border, the SHH locally accumulates, leading to split expression pattern of FGF10. Menshykau *et al.* [37, 26] have proposed a model with additional interactions between SHH and FGF10 that could lead to a Turing-type reaction-diffusion mechanisms for branching morphogenesis: A positive feedback loop is closed if apart from regulation of FGF10 by SHH, FGF10 from the mesenchyme also induces SHH production in the epithelial cells. The model suggests that the SHH ligand-receptor interactions allows the localized spots to stabilize. By letting the growth rate of the tissue domain depend on the level of ligand-receptor signaling, it was shown that the tissue branches out [67]. Similar Turing mechanisms are thought to be at work in the kidney [36]. All in all this work suggests that an intricate signaling network between the epithelium and mesenchyme generates a pattern of growth factors that drives branching by locally upregulating tissue growth.

Alongside the growth factor interaction network discussed above, evidence has accumulated over the last fifteen years that autocrine inhibitory signals, such as TGF-*β* provide a robust mechanism for epithelial branching morphogenesis [44, 50, 63]. The epithelial cells secrete a diffusive signal which, upon binding, inhibits their own proliferation or their own motility. At convexly curved locations the inhibitory signal dissipates more easily than at flat or concave locations, much like heat radiates out more rapidly from a mountain whereas it gets ‘trapped’ within valleys. Using tissue-engineered configurations of cells, Nelson *et al.* [44] have shown that murine mammary epithelial cells exhibit such geometry-dependent sprouting activity. Using an image-based model of murine kidney morphogenesis, the Iber group [1] found that a model in which the epithelium secreted an inhibitory signal more robustly and more accurately predicted the future sites of branching than alternative scenarios, such as the Turing-type system discussed above. The Sneppen team [5] showed that the branching growth of *in vitro* cultures of murine pancreatic cells is well explained using a cell-based model in which the autocrine signal inhibits cell proliferation. These mechanisms are analogous to those based on Laplacian growth proposed previously for branching morphogenesis of cell agglomerates [32, 12, 22, 14]. However, these models based on proliferation do not explain how epithelia can form branching configurations in the absence of growth, as observed in cell cultures [52, 11, 45, 19] and *in vivo* [46, 26].

Here we introduce a hybrid model based on the cellular Potts model (CPM) and partial-differential equations (PDE) to study if such autocrine inhibition of stochastic cell motility suffices for branching morphogenesis. We assume that the local concentration of an autocrine signal inhibits cell protrusion activity at the boundary of the tissue. We first show that this simple mechanism is a sufficient explanation for the in vitro observations by Nelson et al. [44]. Then we show that the same mechanism also suffices to reproduce branching morphogenesis in absence of cross-talk with mesenchymal tissues and in absence of cell proliferation. Finally we study the behavior of the model in presence of proliferation, and show that it suffices to reproduce previously reported behavior of branch epithelia such as self-avoidance.

## 2 Results

*In vitro* observations suggest that cell protrusions are inhibited by the local concentration of TGF-*β* [44], leading to the hypothesis that diffusion of autocrine TGF-*β* drives curvature dependent sites of branching [44]. To test if this mechanism suffices to drive epithelial branching, we developed a hybrid cell based and continuum model (Figure 1). A particularly well-suited modeling framework for this purpose is the cellular Potts model (CPM) [18, 26]. The CPM naturally represents the stochastistic protrusion and retraction of cells as they were observed in mammary epithelial cell cultures [44]. The CPM also naturally represents the collective migration of cells during branch extension. The CPM (Figure 1A) simulates the random motility of cells by mimicking iterative attempts to extend and retract of pseudopods, *e.g.*, filopodia and lamellipodia. We assume that these cellular extensions and retractions are regulated by a chemoinhibitor, *e.g.*, TGF-*β* (Figure 1B). Following Nelson *et al.* [44] we assume that this chemoinhibitor is secreted by the cells, that it diffuses through the extracellular matrix (ECM) in which the cells are embedded, and that it is gradually broken down, *e.g.*, through enzymatic degradation or binding and inactivation in the matrix. The chemoinhibitor inhibits the formation of cellular protrusions in a concentration-dependent manner.

**Figure 1:**
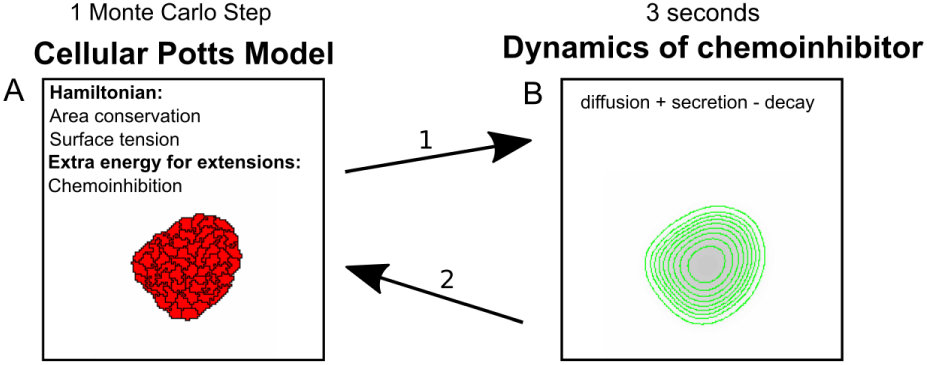
Flowchart of the model. (A) CPM calculates cell movement in tissue due to autocrine inhibition; (B) Autocrine signal is forwarded in space and time, according to PDE in Eq. 4.

### 2.1 Model description

The cellular Potts model describes cells as a collection of lattice sites on a two-dimensional, regular square lattice Λ ⊂ Z^2^. Each lattice site 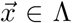 is assigned a spin or cell identifier 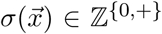, an index of the cell that occupies this lattice site, such that a cell 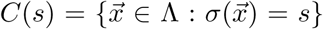, i.e., the set of lattice sites with the same cell identifier *s. C*(0) represents the medium, *i.e.*, the lattices sites 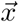 for which 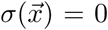. The system evolves by minimizing a Hamiltonian,

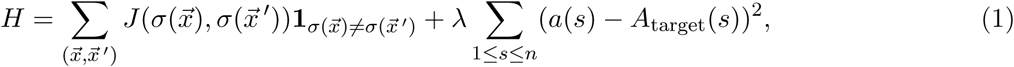

where 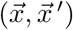 is a pair of adjacent lattice sites, 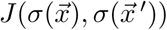 is an interfacial energy, describing cell adhesion and interfacial tensions [30]. The second right-hand-side term introduces a volume constraint, with *a*(*s*) = |*C*(*s*)|, the area of cell *s* and *A*_target_(*s*), the target area of cell *s* and parameter λ a Lagrange multiplier. The CPM simulates cell motility through random attempts to retract or extend the cell boundaries. To simulate a random cell extension or retraction, the algorithm iteratively picks a random lattice site 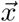, and calculates the energy change Δ*H* resulting from a copy of 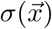 into an adjacent lattice site 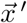. It then accepts this copy depending on the change in energy, Δ*H*, resulting from it, with probability

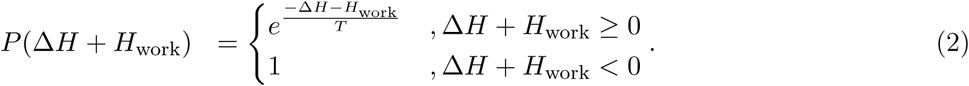

*T* is a motility parameter, aka ‘cellular temperature’, and represents the amount and magnitude of active, random cell fluctuations, which may act against the passive forces given by the energy *H*.

*H*_work_ indicates the energy that is dissipated (*e.g.*, due to friction or viscosity) during a move. It is used here to model chemoinhibition by a field of a secreted chemical, *c*,

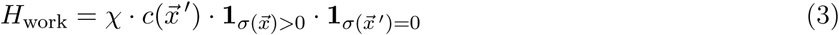

where *χ* regulates the strength of the inhibition, and 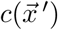 is the chemoinhibitor concentration at the target location 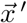. In the cellular Potts model, time is measured in *Monte Carlo Steps (MCS)*, i.e., the number of movement attempts as there are sites in the lattice.

The dynamics of the chemoinhibitory signal is given by a PDE,

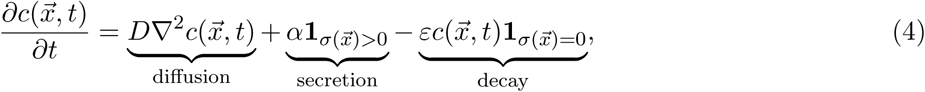

where the secretion and decay terms depend on the current state of the CPM. A simulation consists of consecutive steps of the CPM and the PDE, where one timestep of the CPM is followed by 3 seconds of inhibitor dynamics. This time scale was chosen in accordance to our previous work [39]. The parameter values are given in Table 1.

**Table 1:** Default parameter settings

| Parameter | Description | Value | Unit | Motivation |
| --- | --- | --- | --- | --- |
| $T$ | Cellular temperature | 50 | - | chosen followed by sensitivity analysis (Fig. 6) |
| $A_{\text{target}}$ | Target area | 50 | $\mu\text{m}^2$ | Estimated based on nuclear stainings in Ref. [44] |
| $\chi$ | Chemoinhibition parameter | 25 | - | chosen followed by sensitivity analysis (Fig 5) |
| $\lambda$ | Area constraint strength | 50 | - | chosen |
| $D$ | Diffusion coefficient | $3.75 \cdot 10^{-12}$ | $\mu\text{m}^2 \text{ s}^{-1}$ | chosen together with value of $\epsilon$ based on diffusion length reported in Ref. [44] |
| $\epsilon$ | Decay rate | $5 \cdot 10^{-3}$ | $\text{s}^{-1}$ | Based on half-lives reported in Ref. [64], decay rate in plasma is in range $10^{-4} - 10^{-2}$; diffusion length $\sqrt{D/\epsilon}$ matches observations in Ref. [44] |
| $\alpha$ | Secretion rate | $5 \cdot 10^{-4}$ | $\text{s}^{-1}$ | estimated |
| $J_{01}$ | Cell-ECM adhesion energy | 50 | - | estimated to give slightly adhesive cells |
| $J_{11}$ | Cell-Cell adhesion energy | 20 | - | estimated to give slightly adhesive cells |
| $dt$ | Timestep in PDE integrator | 0.2 | s | numerical stability vs. efficiency |
| $dx$ | Pixel size | $1 \cdot 10^{-6}$ | m | numerical stability vs. efficiency |
| $P$ | PDE iterations per MCS | 15 | - | numerical stability vs. efficiency |
| $t_p$ | Time of increase of area by one lattice site | 250 | MCS | chosen arbitrarily |
| $A_p$ | Area of division | 100 | $\mu\text{m}^2$ | estimated |

### 2.2 Model mimics experimental observation of branching at convex sites

Nelson and coworkers [44] have reported mammary epithelial cells show geometry-dependent sprouting. They cultured murine mammary epithelial cells inside small micropatterned cavities stamped into collagen gels. After induction with growth factors the cells formed multicellular sprouts preferentially at the positively curved (convex) parts of the cell clusters (Figures 2A and B). At crevices between two cell clusters, where secrete growth factors accumulate, no sprouts formed (Figures 2C and D). To test if our mathematical model suffices for explaining these observations, we initialized our model simulations with the shapes used in the experiments by Nelson *et al.*[44] *(Figures 2 E-H). The size of the geometry matched those used in the in vitro* experiments [44] and we used the parameters in Table 1, for which there are on average 5 cells across the diameter, corresponding to nuclei counts in Ref. [44].

**Figure 2:**
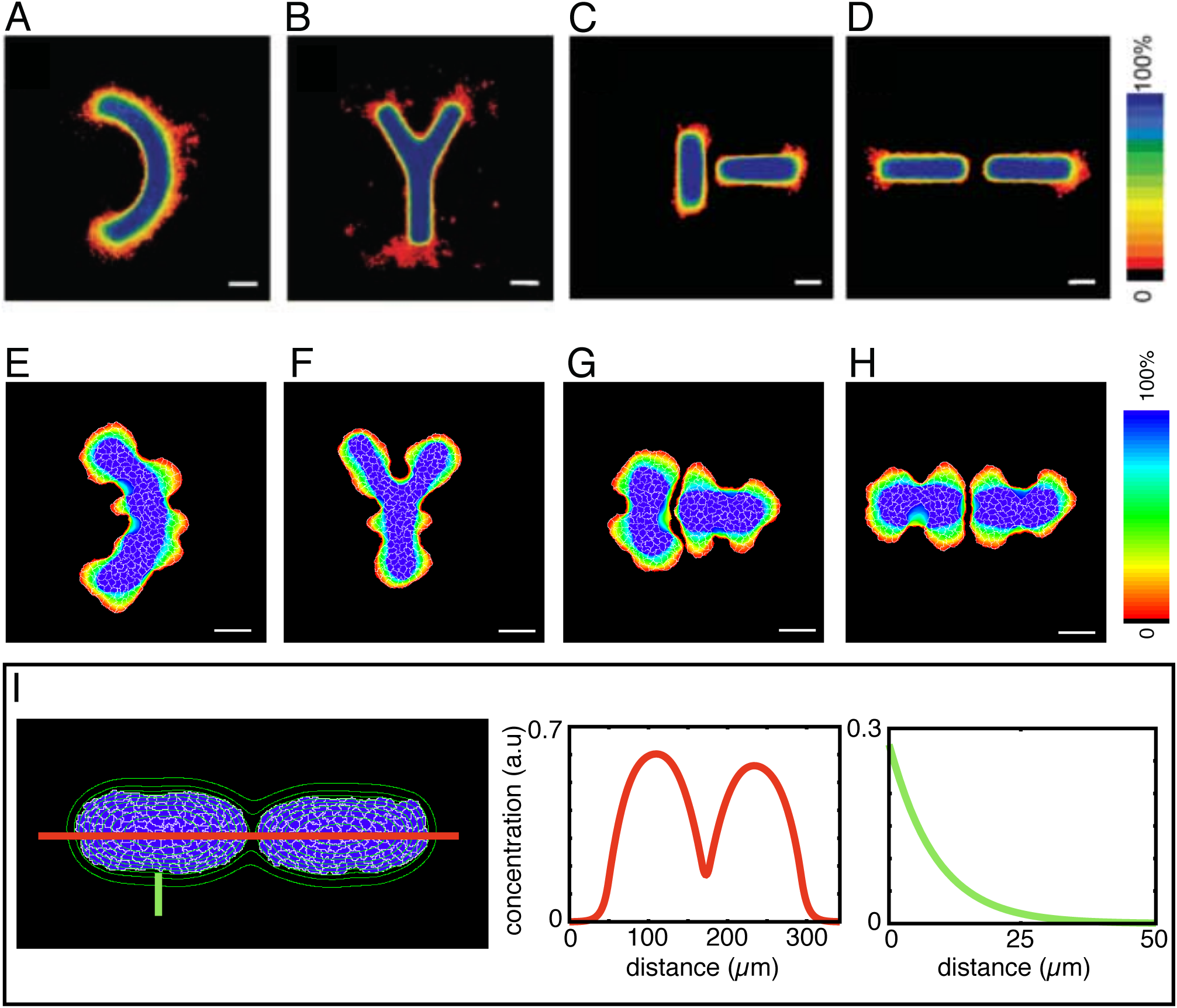
Model replicates branching at convex sites of geometries. Heat map indicates the percentage of time a cell was present at a given site during 4000 Monte Carlo Steps. (A) Experimental frequency map of epithelial cells for the reversed “C” shape; (B) Experimental frequency map of epithelial cells for the “Y” shape; (C) Experimental frequency map of epithelial cells in two aligned rectangles; (D) Experimental frequency map of epithelial cells in two orthogonally placed rectangles; (E) Simulated cells placed in a reversed “C” shape - see also Video S1; (F) Simulated cells placed in a “Y” shape - see also Video S2; (G) Simulated cells placed in two aligned rectangles - see also Video S3; (H) Simulated cells placed in two orthogonally placed rectangles - see also Video S4; (I) Concentration profiles of the inhibitory, autocrine signal for the simulation shown in panel H. Panels A-D reprinted from Ref. [44] with written permission from the AAAS (#4680251276893). Parameter values for the simulations as in Table 1. Scale bars: 50 µ*m*

We first ran the CPM model for 750 MCS during which time cell shapes could equilibrate. For the next 750 MCS we simulated only the PDE such that the chemical field had reached a steady-state (Figure 2I). The shapes and lengths of these simulation steady-state gradients matched well with the experimentally-observed TGF-*β*1-gradients (compare Figure 4A in Ref. [44] with Figure 2I). We then simulated the CPM and the PDE concurrently for a further 4000 MCS, allowing for some cell expansion (1 *µm*^2^ of additional target area per 100 MCS), but disabling cell division. Frequency maps show the percentage of the last 4000 MCS that a cell is present at the site. Similar to the *in vitro* observations [44], *in silico* multicellular sprouts appeared preferentially at the convex parts of the geometry. For the “C” shape, branching occurs at the tips and the “belly” of the “C”, which are the convex parts (Figure 2E and Video S1). Similarly, for the “Y” shape, sprouts appeared around the convex tips (Figure 2F and Video S2). Our model could also reproduce the setup shown in Figure 2(C,D). Here cells were placed into two rectangular wells positioned at 90^◦^ angles or adjacent to each other. As in the C and Y-shaped geometries, the cells sprouted at the convex regions of the morphologies away from the other morphologies (Figures 2(G,H) and Videos S3 and S4). No sprouts were formed at the convex regions near the crevice between the two geometries due accumulation of the inhibitory signal (red curve in Figure 2I). This observation is correctly reproduced in the simulations. A discrepancy between the predictions of our simulations and the experimental observations is that we needed to keep a bit of pressure on the cell boundaries through a slow cell expansion term. In absence of this term, the concave cell boundaries moved inwards as can still be observed in Figure 2(G,H); a more refined representation in the CPM of the ‘cavities’ that contained the cells in the experiments will likely prevent this inward motion and hence improve the predictions. Possibly this also solves a further discrepancy with the experiments: the cell expansion term generates sites of cell protrusion that are not seen in the experiments: see, *e.g.*, the lateral protrusions adjacent to the crevice in Figure 2H and the lateral protrusion below the ‘fork’ of the Y in Figure 2F.

**Figure 3:**
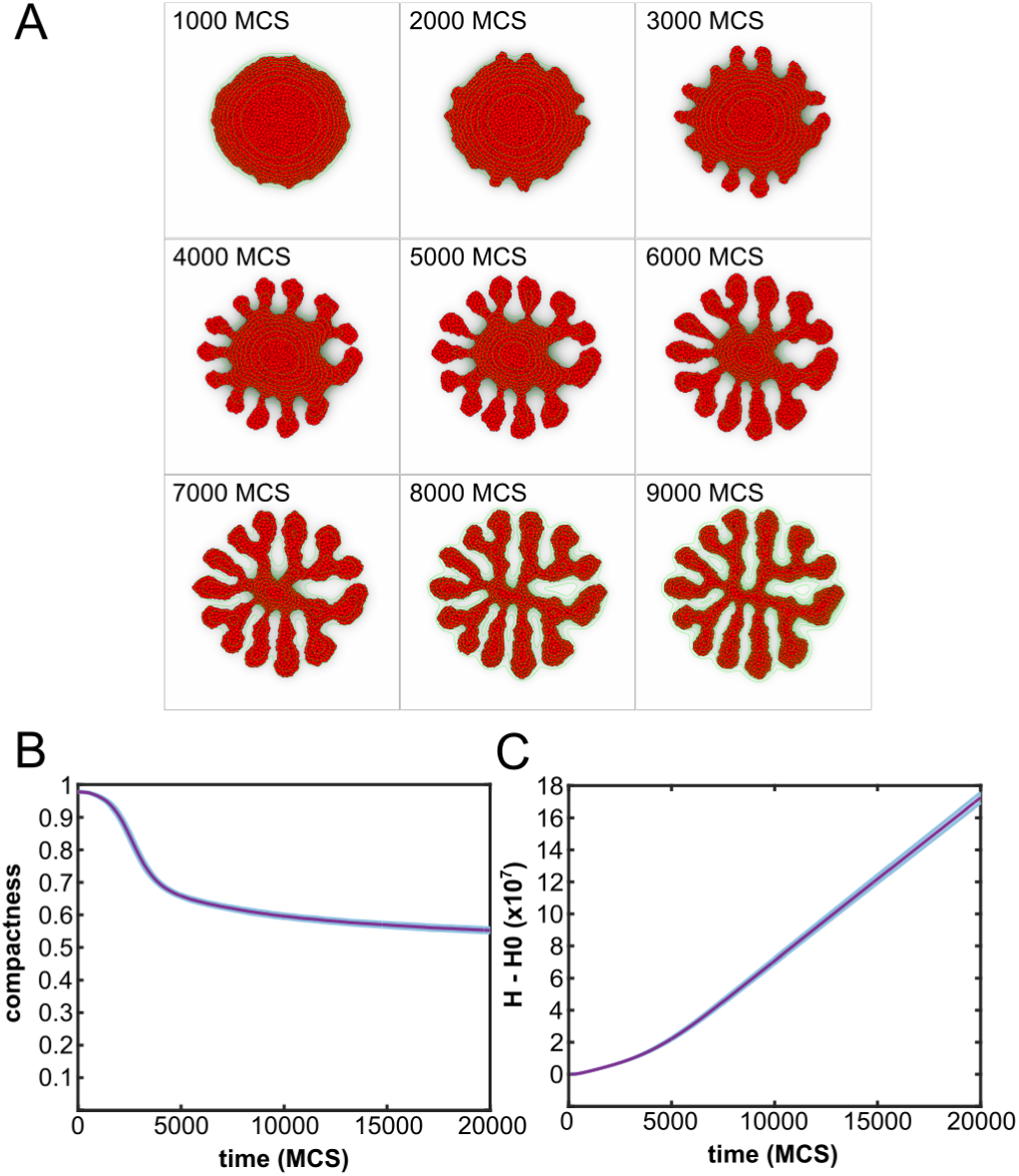
Simulation of branching by autocrine inhibition. (A) Timelapse of a model realization - see also Video S5; (B) Compactness as a function of time, shaded regions: standard deviations of 100 simulations; (C) Energy spend by the system (*H* − *H*_0_ = Σ Δ*H*) as a function of time, shaded regions: standard deviations of 100 simulations.

**Figure 4:**
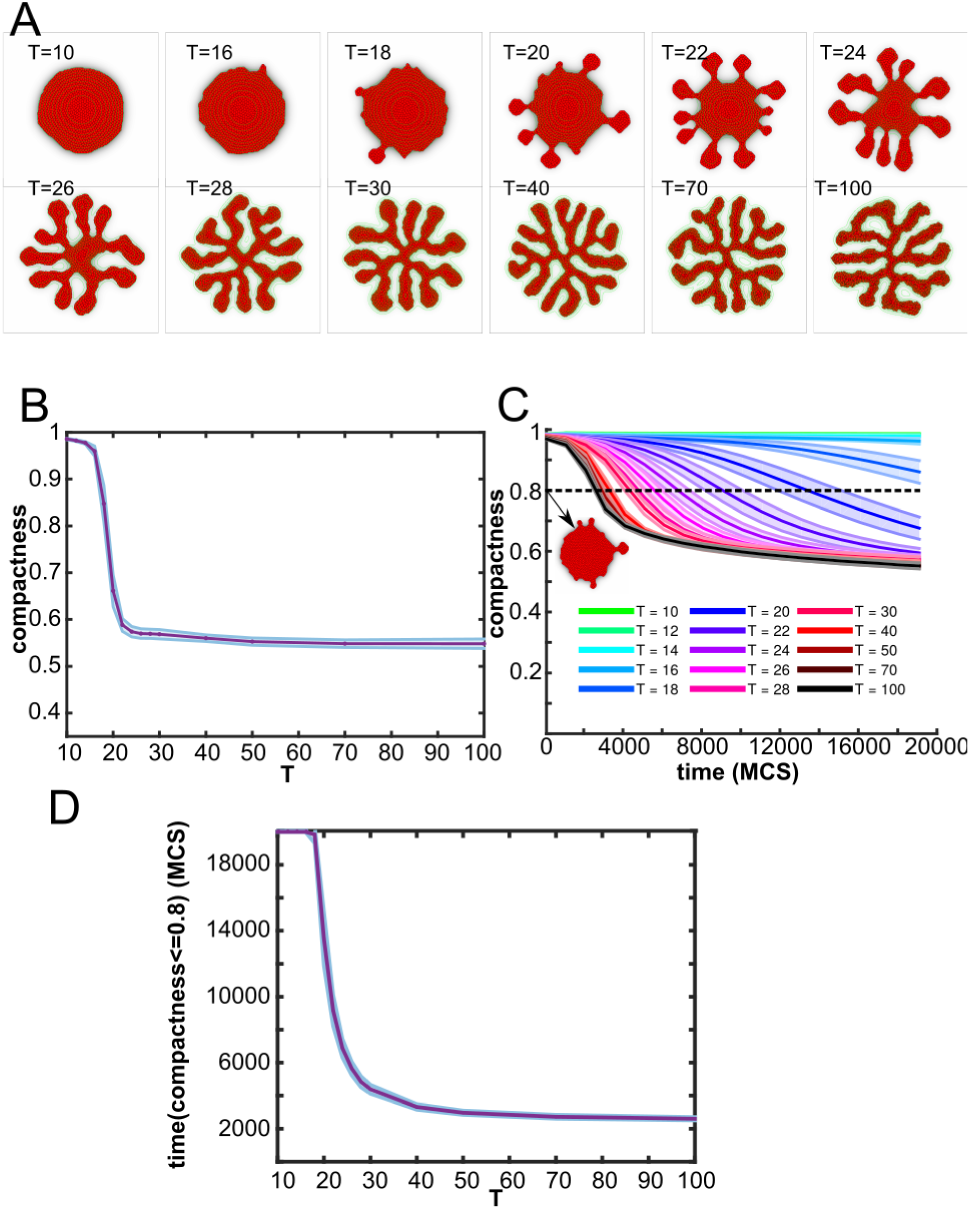
Cellular temperature regulates branching dynamics. (A) Example configurations of the tissue at 20000 MCS for different values of *T* ; (B) Compactness as a function of *T*, shaded regions: standard deviations of 100 simulations; (C) Compactness as a function of time, shaded regions: standard deviations of 100 simulations; different colors: different values for *T*, see legend.

### 2.3 Autocrine inhibition of cell movement drives branching

We next asked if this mechanism proposed in Ref. [44] also suffices for branching morphogenesis. We initiated the model simulation with a disc-shaped structure radius 0.225 mm in a lattice of 0.9 mm by 0.9 mm containing approximately 1000 cells. Figure 3A and Video S5 shows a model simulation for the first 9000 MCS. A first look at the time series of the simulation shows that after approximately 1000 MCS the boundary of the disc becomes bumpy. Then around 3000 MCS, many droplet-like extensions appear. The length of these extensions increase and as a result, a fully branched structure, with evenly thick branches is formed, that stabilizes around 8000 MCS. To test if and to what extent this result depends on the numerical resolution and scaling of the simulation (Δ*x*) we have repeated it for Δ*x* = 5 · 10^−7^*m* (Video S6) and for Δ*x* = 2.5 · 10^−7^*m* (Video S7), *i.e.* twice and four times the original resolution. At these refined resolutions the simulations progressed more slowly due to the reduced length scale of the cellular extensions and retractions, but otherwise the results did not depend on the spatial resolution. We thus performed our parameter studies at Δ*x* = 1 · 10^−6^*m* for computational efficiency.

To quantify branching, we define the compactness of the morphology as *C* = *A*_tissue_*/A*_hull_ [39], *i.e.*, the ratio between the area of the largest connected component of the tissue and the area of its convex hull, the smallest convex polygon that contains the tissue [17]. A compactness of 1 implies a perfectly circular tissue shape, whereas a low value of the compactness indicated a high degree of branching or high degree of cell scattering. The compactness of the morphology rapidly drops during the first 5000 MCS and slowly decreases after that (Figure 3B) indicating slow thinning of the branches.

Initially the boundary of the circular tissue roughens due to random cell motility. While the secreted inhibitor accumulates at the concave locations of the morphology, it diffuses away more easily at the convex locations (Figure 3A). As a result, cell motility is inhibited more strongly at the concave and flat regions than at the convex regions of the morphology, such as the branch tips. Thus the secreted inhibitor leads to a geometry-dependent rate of cell extension. Cell motility is strongly inhibited at the ‘valleys’ between the branches, but is more frequent at the sprout tips. This leads to a ‘ratchet’-type, dissipative branching mechanism: cells attempt to extend and retract randomly, and at sufficiently high temperatures they can do so even against the local energy gradient generated by the inhibitor (Eq. 2). Due to the secreted inhibitor, such extensions against the energy gradients are more frequent at branch tips than in the ‘valleys’.

To test if indeed branching is due to a ‘ratchet’-like, dissipative mechanism, we measured the cumulative energy of the system 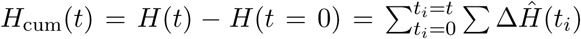. Indeed it increases as a function of time (Figure 3C), showing that for many of the moves Δ*H* > 0 holds, acting against the large positive energy contribution *H*_work_ due to inhibitor of cell motility (Eq. 4).

### 2.4 Random motility regulates branch initiation and branching speed

The previous section showed that the proposed branching mechanism is driven by random cell motility. Interestingly, Btbd7, a positive regulator of epithelial cell motility, is required for branching morphogenesis of salivary glands and the lung [48], and is expressed in branching end buds in a variety of branching epithelial organs [6]. Knock-out of Btbd7 reduces epithelial cell motility in the end buds of submandibular salivary glands, leading to incomplete branching [6]. We thus hypothesized that the level of random cell motility should have similar effects in our mathematical model. The distribution of the magnitudes of the random forces exerted by the cell extensions and retractions is given by parameter *T*, the motility parameter aka cellular temperature (Eq. 2). Figure 4A shows morphologies for increasing values of *T*. Consistent with the inhibition or knockout of Btbd7 [48, 26] at low values of *T* the tissue did not branch, because only few random cell protrusions were strong enough to overcome the effect of the chemoinhibitor. For slightly higher cell motility, at cellular temperatures of around *T* = 20, the tissue developed droplet-like extensions: as soon as one or a few protrusions locally overcame the effect of the inhibitor the curvature locally increased leading to reduced levels of chemoinhibition. For elevated values of *T* the tissue branched normally and the branches became longer and thinner than for lower values of the cellular temperature.

Indeed the compactness of the morphologies formed after 20000 MCS declines sharply for increasing values of *T* up to around *T* = 20 (Figure 4B), reflecting that branching occurs from around *T* = 20. Figure 4C shows the compactness as a function of time for increasing values of *T*. For low values of *T* (*T* = 18 and *T* = 20) the compactness decreased slowly over time, but it did not reach a low compactness before the end of the simulation, while for higher values of *T* branching accelerates. We quantified speed of branching by measuring *t*(*C* = 0.8), the time required for the tissue to drop below a compactness 0.8 (dashed line in Figure 4C). Figure 4D shows that the speed of branching quickly increases with the value of *T* and then saturates. In conclusion, consistent with experimental observation, the motility parameter *T* regulates the initiation and the speed of branching and has some effect on the branching morphology.

### 2.5 Strength of autocrine inhibition has a biphasic effect on branching

The previous section showed how the cellular temperature affected branching dynamics. We next studied the effect of the chemoinhibition strength *χ*. Because the value of *χ* and *T* both determine the probability that a cellular protrusion is accepted (see Eqs. 2 and 3), the effect of *χ* and *T* are likely interrelated. Interestingly, the chemoinhibition strength (Figure 5A-B) has a biphasic effect on branching. For relatively low values of *χ* = 50, the morphology retained its circular shape and no branches formed or they remained very short (Figure 5C) (the increased values of the branch length for low values of *χ* in Figure 5C are due to artifacts of the skeletonization algorithm). For these low values of *χ* the impact of the autocrine signal is negligible, such that the dynamics will dominated by the ‘standard’ Hamiltonian (Eq. 1). At higher values of *χ* the differences in the levels of the autocrine signal concentration around concave and convex regions differ sufficiently for the curvature-effect to set in and branch formation to occur, as shown by the reduced compactness (Figure 5B) and increased branch length (Figure 5C). At even higher values of *χ*, the branches become thinner and longer. As the chemoinhibition is active only at cell-ECM interfaces, at higher values of *χ* the chemoinhibition term becomes dominant over the other components of the Hamiltonian including the surface tension. This also allows droplets to break off of the spheroid; this is an irreversible process, because the chemoinhibitory field makes it energetically very costly for such droplets to join the morphology again. For the largest values of *χ* tested, branches did not form, because cell protrusions became very costly. In conclusion, the chemoinhibition strength *χ* regulates the degree of branching in a biphasic manner.

**Figure 5:**
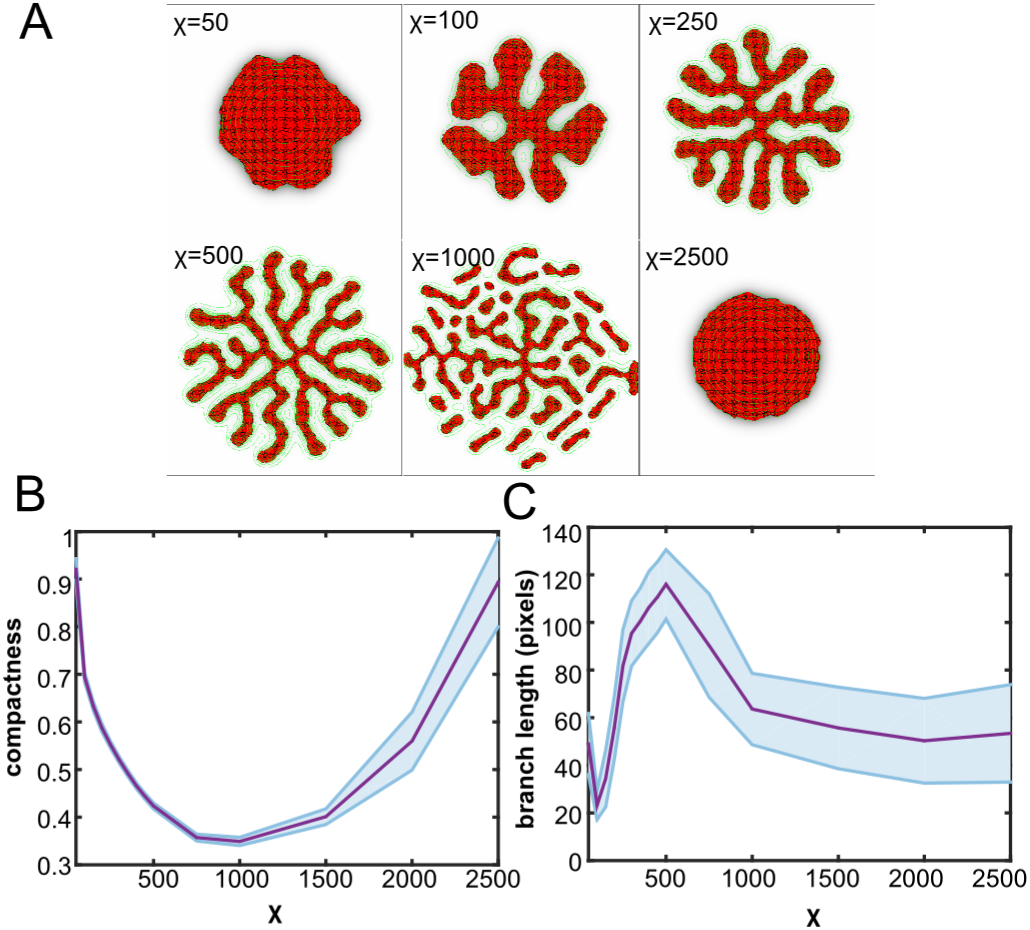
Strength of autocrine chemoinhibition biphasically drives branching. (A) Example configurations of the tissue at 20000 MCS for different values of *χ*; (B) Compactness as a function of *χ*, shaded regions: standard deviations of 100 simulations; (C) Branch length as a function of *χ*, shaded regions: standard deviations of 100 simulations.

### 2.6 Decreasing surface tension promotes branching

In a study where lung epithelium was isolated *in vitro*, Hartmann and Miura showed that a decrease in surface tension, by disruption of the cytoskeleton using cytochalasin D, results in more but smaller branches [22]. Similarly, inhibiting cell contractility, thus reducing surface tension, promotes branching morphogenesis in pancreas [19]. We therefore studied how the value of of surface tension affected branching morphogenesis. In the CPM, surface tension *γ*_01_ is defined as 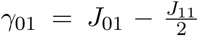 [16], where *J*_01_ is the interfacial energy between cells and medium and *J*_11_ is the interfacial energy between two cells. Figure 6 shows the effect of *γ*_01_ on branching morphogenesis. In agreement with the experimental data, increased surface tension reduces the number of branches and gives thicker branches. For very low values of *γ*_01_ there are many thin branches, which occasionally merge. In simulations with increased surface tension the branches became thicker (Figure 6B) and shorter (Figure 6C). For the highest values of *γ*_01_ tested, the branches became droplet-shaped and had more variable thickness than for lower values of *γ*_01_. In agreement with [22] these results illustrate that the surface tension *γ*_01_ acts as a restoring force that counteracts the curvature effect due to chemoinhibition.

**Figure 6:**
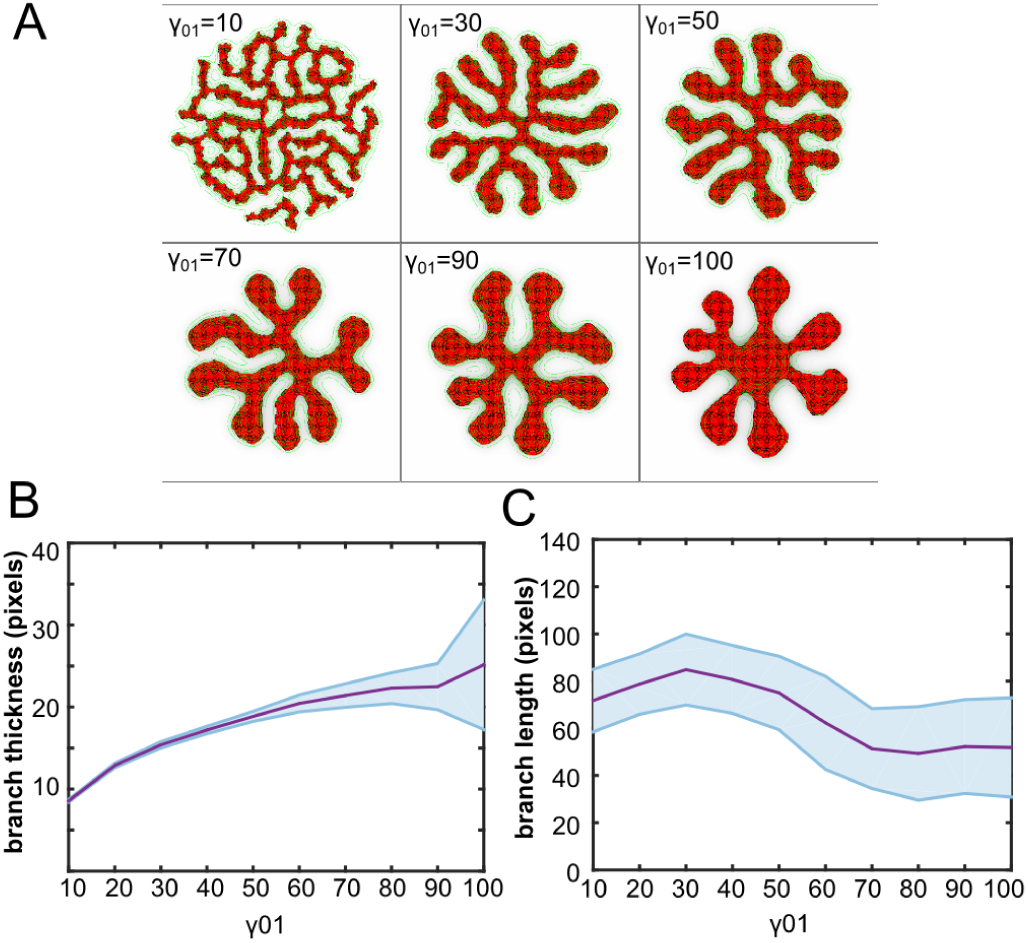
Surface tension affects branch morphology. (A) Example configurations of the tissue at 20000 MCS for different values of *J*_01_; (B) Branch thickness as a function of *γ*_01_, shaded regions: standard deviations of 100 simulations; (C) Branch length as a function of *γ*_01_, shaded regions: standard deviations of 100 simulations.

### 2.7 Space-filling branching growth and branch avoidance in presence of cell proliferation

The results above suggest that autocrine inhibition of random cell protrusion is sufficient for branching morphogenesis of epithelial cells, even in absence of cell proliferation and in absence of regulatory interactions with the mesenchyme. However, *in vivo* branching morphogenesis usually requires cell proliferation [11]. Here we investigate, therefore, how the proposed branching morphogenesis mechanism behaves in the presence of cell proliferation. To mimic cell proliferation, the target area was incremented by 1 once every 100 MCS. Cells divided over their short axis after the actual area had reached a threshold value; thus pressure from adjacent cells inhibited proliferation.

Figure 7 shows a simulation initiated from a circular blob of proliferative cells. The insets in the final configuration highlight two proliferation events (purple and cyan dots). With proliferation, the mechanism produces a space-filling branching structure. The branches did not merge: branch tips that grew towards each other were repelled by one another (Figure 7; black circle) or they are terminated. Such ‘self-loathing’ [31] of branches is due to the accumulation of the autocrine inhibitor between branches (Fig. 2G, H) and has been observed in ex vivo cultures of murine urinary collecting ducts [9] and mammary gland tissue [21]. Note that in our model, the auto-inhibition is responsible for the branching morphogenesis itself, and also gives rise to ‘self-loathing’ [31] of adjacent branches, leading to branch avoidance.

**Figure 7:**
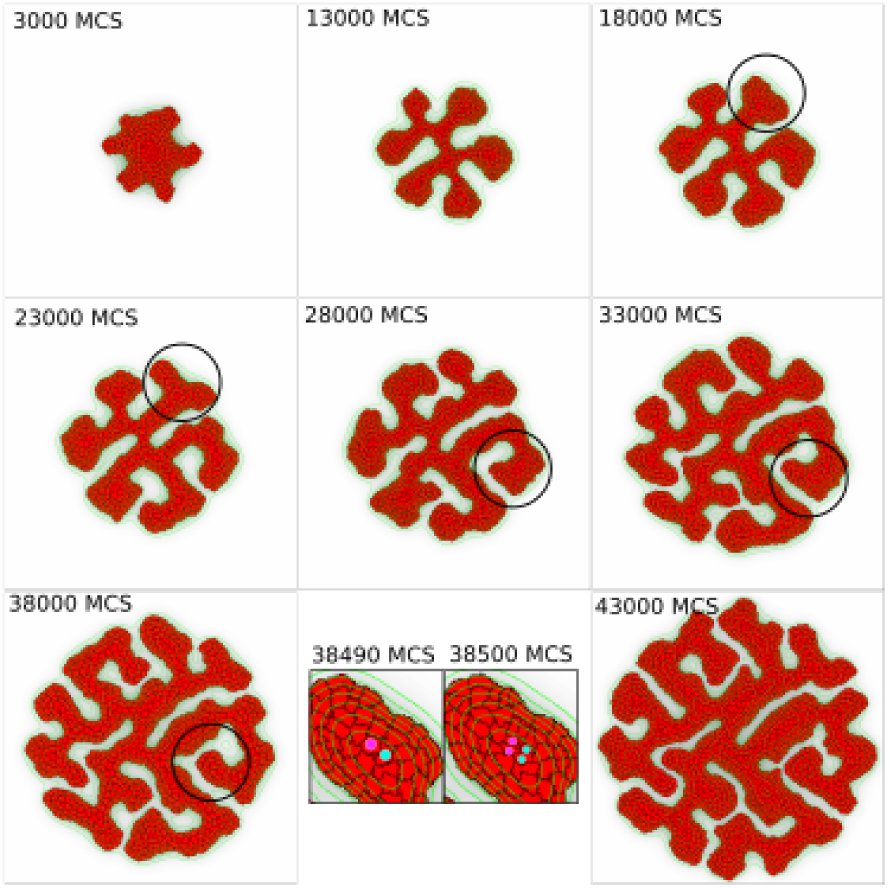
Branching with cell proliferation. Timelapse of a model realization. The black circle tracks a branch avoidance event. The inset at 15500 MCS shows two cell division events. All parameter values are given in Supplementary Table S3.

## 3 Discussion

Using mathematical modeling we have shown that the mechanism for geometry-dependent sprouting in tissue-engineered constructs of mammary epithelial cells proposed by Nelson and coworkers [44] suffices for autonomous branching morphogenesis of epithelial tissues, in absence of cell proliferation and interaction with the surrounding mesenchymal tissues. Importantly, and in contrast to related models based on Laplacian growth principles, the present model produces sprouts and branching structures in absence of proliferation. The model suggests that branching morphogenesis can occur due to a ‘ratcheting’ mechanism, which favors random cellular protrusions at convex locations of the morphology over protrusions at concave locations. The present model derives conceptually and methodologically from our previous work on angiogenesis [39], in particular from the model variant based on ‘extension-only chemotaxis’ (see Section ‘A Dissipative Sprouting Mechanism’ in Ref. [39]). The key difference with this previous work is that the autocrine *chemoinhibition* mechanism studied here is based on the *concentration* of 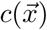 (Eq. 3), whereas the chemotactic mechanisms studied previously relied on chemical gradients 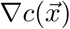. This small difference has a large effect on the patterns predicted by the model: the present mechanism explains the formation of self-avoiding branching patterns, as observed in many epithelial-derived branching organs, whereas the previous work predicted the formation of network-like patterns, such as those found in microvasculature. In the presence of proliferation our model is similar to previous models based on Laplacian growth. In these models, tissues branch due to a Mullins-Sekerka instability, where positive curvatures experience higher gradients of a pressure field [32], a growth-promoting [22] or a growth-inhibiting field [5]. In particular the latter model by the Sneppen group [5], which is based on growth-inhibition resembles ours. However, our model differs from it in an essential aspect: our model can explain the branching morphogenesis in absence of proliferation.

The assumptions of the model are simple, but plausibly based on experimental observation [44], and its predictions agree surprisingly well with some observations of branching epithelial organs. At the same time, there are of course many observations that our model cannot explain and that will open up new perspectives for future modeling studies. In particular, observations of renal epithelial cells challenge the hypothesis of autocrine inhibition of motility studied in this work [35]. Renal epithelial cells grown on micropatterns of specific curved geometry exhibit curvature dependent protrusions. These are potentially regulated by autocrinically secreted BMP7, a member of the TGF-*β* superfamily. To test this hypothesis, Martin *et al.* [35] applied a flow to the culture medium that should flush away any diffusive signals potentially secreted by the cells. Surprisingly, this treatment did not affect the curvature-dependent protrusions. The authors suggested that membrane tension might be responsible for the curvature effect instead. Pavlovich et al. [50] argued that the autocrine inhibition mechanism may uniquely apply to mammary epithelial patterning. An alternative explanation for the observations by Martin *et al.* [35] may be that autocrinically secreted signals are bound to secreted extracellular matrix (ECM) proteins, which would prevent them from being flushed away. Such pericellular retention of signaling molecules has been observed for VEGF in endothelial cells [29]. Indeed the activity of TGF-*β* activity is likely regulated by the chemistry and the mechanics of the ECM. TGF-*β* is bound to the ECM in a latent form. The active moiety can be released from the ECM through proteolysis [59] or through mechanical stretching of TGF-*β* [23]. Further supporting the importance of the ECM in branching morphogenesis, reducing the cytoskeletal tension (likely reducing the cellular traction forces exerted on the ECM) reduces the number of branches formed in embryonic lung explants [42, 26]. In our previous work we have modeled how TGF-*β* release from fibrin matrices can regulate angiogenic sprouting [4], and how mechanical cell-ECM interactions can coordinate pattern formation and sprouting in endothelial cell cultures [60, 26]. In our ongoing studies we are incorporating these two approaches into our models of epithelial branching. These model extensions may provide deeper insight into the role of the ECM in branching morphogenesis. Apart from the leads provided by the observations of Martin *et al.* [35], another direction for further research may consider TGF-*β* release through ECM proteolysis and mechanical straining of the ECM by the epithelial cells. Possibly such models of the mechanochemical cross-talks in TGF-*β* signaling can help reconcile some of the experimental observations on the role of membrane tension and mechanical force generation in branching morphogenesis.

Another lead that may provide new insights into branching morphogenesis concerns the analogy with scratch assays for wound healing. In such assays, epithelial cells migrate into a free surface left open by the scratch [51, 26]. During closing of the scratch the epithelial cells organize into finger-like structures, which will eventually merge to close the free space. The finger-like structures extending from the boundary look similar to initial branching structures, and like in branching, proliferation does not seem to be the driving factor in fingering experiments [51] nor in mathematical models of fingering [49, 26]. With a cellular Potts model, Ouaknin *et al.* [49] showed that if cells in contact with free space secrete a signal to which all cells chemotact, fingering into the open space occurs. In this model, cells at the boundary move and drag cells along, which will encounter more open space which increases the chemotactic signal and thus reinforces fingering. In a model by Mark *et al.* [34], the Mullins-Sekerka instability arises from a curvature dependent cell motility. In a follow-up paper which included velocity alignment of cells [58], it was shown that the thickness of the fingers depends on the length scale on which the cells in the bulk align their velocities. Indeed, epithelial cells in fingers are observed to move together in a highly coordinated fashion [53]. The relation between motility and curvature was based on observations where epithelial cells have a higher protrusion rate at convexely curved surfaces [55].

For computational efficiency, the analyses in this work are based on two-dimensional simulations. However, it is straightforward to extend the model to three dimensions (*cf.* Video S1 for an example with proliferation). Our two-dimensional simulations represent quasi-2D cultures, *e.g.*, mammary epithelial in thin fibrin gels [41] or kidney rudiments cultured on filters supported by Trowell screens [9]. To represent such quasi-2D situations, we have assumed that the decay of the autocrine signal only takes place outside of the cells and the signal only affects cell motility at the periphery. An alternative interpretation of a two-dimensional model could be the projection of the three-dimensional case in two dimensions. In this interpretation of the two-dimensional model, we should also consider the degradation of the signal underneath the cells.

The present model is of course a great simplification of epithelial branching morphogenesis *in vivo*. Epithelial morphogenesis involves interactions of many signaling molecules and receptors from the epithelium and mesenchyme and, despite some similarities, there are large differences between organs of epithelial origin. The model generates variable branching patterns such as in mammary epithelial tissues [56, 26], whereas other organs such as lung and kidney display highly stereotypic, reproducible patterns of branching [40, 26]. Future work could explore the hypothesis that additional signaling molecules and the interaction with the surrounding mesenchyme and the ECM could fine-tune generic branching mechanisms such as the one presented here, leading to more stereotypical branching.

## 4 Methods

### 4.1 Model implementation

The model was implemented using the Tissue Simulation Toolkit (http://sourceforge.net/projects/tst). The PDE is solved by using a forward Euler method on a regular square lattice matching that of the CPM. *i.e.*, Δ*x* = 2*µ*m. The CPM and PDE are coupled using an operator splitting approach: After running the Cellular Potts model for one Monte Carlo step, 15 of the numerical integration steps are performed with Δ*t* = 0.2 seconds, such that the PDE runs for *t*_*c*_ = 3 seconds per MCS. As initial conditions for the PDE we assume 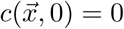 for all 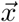. We used zero boundary conditions, *i.e.,* 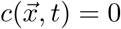 at all boundaries.

### 4.2 Morphological measures

The morphologies were characterized using the following morphometric measures:

#### 4.2.1 Compactness

The compactness is defined as the ratio between *A*, the domain covered by the tissue, and the area of its convex hull *A*_hull_ [17]:

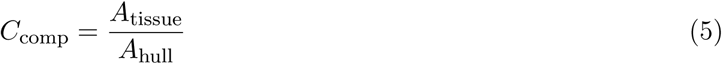

The convex hull is the smallest convex polygon that encloses the object of interest. A compactness of *C*_comp_ = 1 indicates a convex tissue shape, whereas a lower value of the compactness indicates branching or cell scattering.

#### 4.2.2 Branch length

In order to find the branches of the structure, we generate the morphological skeleton of the tissue [20, 26]. Using this skeleton image, we calculate the length of the branches as follows. For every edge, the two nodes of the edge are removed from the skeleton image by removing all lattice sites around the nodes with increasing radius, until a radius *w* is found such that the skeleton image is divided in at least two separate components, of which one is the edge of interest. The length of branch is then determined by counting the pixels that make up the branch and adding twice the radius *w* to the final result.

#### 4.2.3 Branch thickness

To calculate the branch thickness, we adopted an approach by Filatov *et al.* [13]. *We take a* to be the image of the tissue and let *b*(*r*) be a disk *b*(*r*) = {(*x, y*) ∈ R^2^ : *x*^2^ + *y*^2^ ≤ *r*} with variable radius *r*. The branch thickness can now be defined as the value of *r* for which branches disappear out of the morphological opening *a* ◦ *b*(*r*).

The area of the morphological opening decreases with the radius *r* (Figure 8), because more branches disappear from the image with increasing *r*. We approximate the branch thickness by finding a point where this graph decreases sufficiently fast and then becomes flat, indicating that most branches have disappeared. At some point the graph becomes more or less horizontal. This region corresponds to the circular part of the tissue, in which many circles *b*(*r*) fit. So, the value for *r* for which the graph becomes horizontal indicates is the thickness of the branches. We detect this horizontal region by first finding a region where the graph decreases sufficiently fast and then a region where it decreases slowly. Let *M*_*A*_(*r*) be the area of *a* ◦ *b*(*r*).

**Figure 8:**
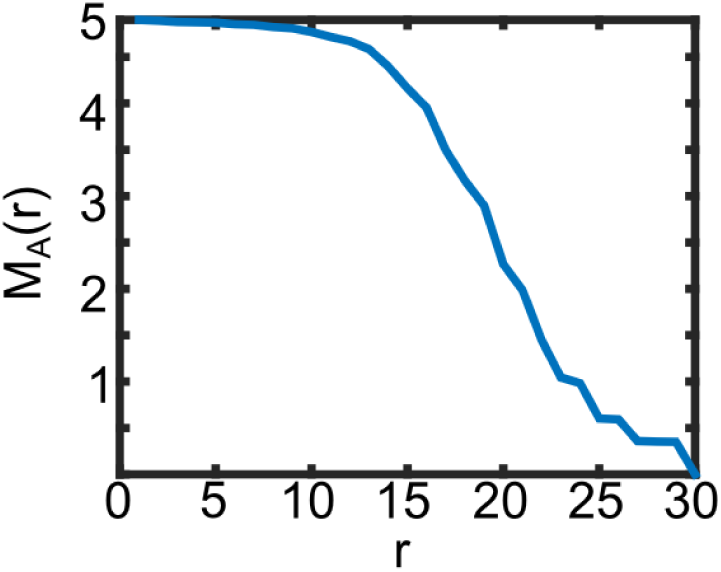
Example of the radius. *r* **of a branching morphology as a function of the area of the morphological opening** *a* ◦ *b*(*r*).

We find an 0 < *r*_1_ ≤ *r*_max_ for which *M*_*A*_(*r*_1_) − *M*_*A*_(*r*_1_ − 1) < *a*_1_ and then the value *r*_1_ < *r*_2_ ≤ *r*_max_ for which *M*_*A*_(*r*_2_) − *M*_*A*_(*r*_2_ − 1) > *a*_2_ (*r*_2_ is set to *r*_max_ if such a value does not exist). The branch thickness is then found by taking the value of *r* for which *M*_*A*_(*r*) is closest to 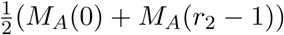. The values of *a*_1_ and *a*_2_ are determined empirically. The value of *r*_max_ is set to 30 to reduce computation time.

In case no branches or only very small branches are present (*M*_*A*_(*r*_1_) − *M*_*A*_(*r*_1_ − 1) ≥ *a*_1_ for all 0 < *r*_1_ ≤ *r*_max_) we apply the following algorithm. When the decrease in *M*_*A*_(*r*) is not larger than *a*_1_ in the entire graph we simply take the distance from the center of mass of the tissue to an ECM point in four different directions and select the lowest distance as the radius. In this case, the radius represents the radius of the unbranched cell aggregate but we take this as the branch thickness. We follow this approach, because increasing the radius to the width of the initial circular tissue (typically more than twice as large as *r*_max_ = 30) and repeatedly computing *M*_*A*_(*r*) would require excessive computation time.

## 5 Supplementary Videos

**Video S1** Video of the simulation shown in Figure 2E. Length: 5000 MCS; frame rate: 100 MCS per second.

**Video S2** Video of the simulation shown in Figure 2F. Length: 5000 MCS; frame rate: 100 MCS per second.

**Video S3** Video of the simulation shown in Figure 2G. Length: 5000 MCS; frame rate: 100 MCS per second.

**Video S4** Video of the simulation shown in Figure 2H. Length: 5000 MCS; frame rate: 100 MCS per second.

**Video S5** Simulation of branching by autocrine inhibition of cell motility, corresponding to Figure 3A. Length: 8500 MCS; frame rate 100 MCS per second.

**Video S6** Simulation of branching by autocrine inhibition of cell motility at increased resolution Δ*x* =5 · 10^−7^*m*; all other parameters as in Figure 3A. Length: 20000 MCS; frame rate 1000 MCS per second.

**Video S7** Simulation of branching by autocrine inhibition of cell motility at increased resolution Δ*x* = 2.5 · 10^−7^*m*; all other parameters as in Figure 3A. Length: 20000 MCS; frame rate 1000 MCS per second.

**Video S8** Simulation of branching growth by autocrine inhibition of cell motility and with cell proliferation, corresponding to Figure 7. Length: 45000 MCS; frame rate 1000 MCS per second.

**Video S9** Example of three-dimensional simulation of the autocrine inhibition model in the presence of cell proliferation.

